# Human lower leg muscle pump acts like a stream diversion pump during locomotion

**DOI:** 10.1101/2023.09.25.559436

**Authors:** Roman A. Tauraginskii, Fedor Lurie, Sergei Simakov, Rishal Agalarov, Pavel Khramtsov, Maxim Babushkin, Denis Borsuk, Maxim Galchenko

## Abstract

**Background:** Calf muscle pump (CMP) failure is associated with the development and progression of chronic venous insufficiency as characterized by ambulatory venous hypertension (AVH). However, the explicit interconnection between AVH with CMP failure remains uncertain because the concept of CMP function is controversial. The study aimed to measure pressure in different segments of the great saphenous vein (GSV) and intramuscular vein of the gastrocnemius (GCM) during exercise.

**Methods:** Twelve legs of nine healthy volunteers were enrolled in the study. Continuous pressure (IV_CALF_, GSV at ankle, proximal calf, and mid-distal thigh) and electromyography data (GCM and anterior tibial muscle [ATM]) were recorded during treadmill walking, running, and plantar flexion exercises. The pressure gradient (PG, mmHg) between adjacent points of measurement was calculated. Minute unit power of muscle pump ejection and suction (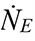, and 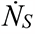, MPa/min) were calculated and compared with the arterial blood supply of the lower extremity (LBF, L/min).

**Results:** PG demonstrated a consistent pattern of changes during walking and running. An absence of PG directed from the calf to the thigh (centripetal) in the GSV was observed. Instead, a retrograde PG was verified throughout the entire stride cycle. Its value decreased with the increase in stride cycle frequency. The dynamics of PG between the IV and GSV were the following: It was directed from the IV to GSV during GCM contraction and was reversed during ATM contraction and GCM relaxation (swing phase). LBF, 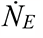, and 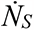 demonstrated similar exponential growth with the increase in stride frequency during walking and running.

**Conclusions:** The pressure gradient in the GSV prevents centripetal flow during locomotion. Instead, PG directs blood flow from the GSV toward intramuscular veins. The increase in CMP unit power was tightly coupled with lower extremity arterial blood supply growth that prevented ambulatory venous pressure rise during exercise.

**Novelty and Significance What Is Known?:**

- The existence of a pressure gradient in superficial veins directed centripetally (from calf to thigh level) has been confirmed for the state of rest only (lying and standing).
- The existence of a pressure gradient directed from intramuscular to superficial veins or vice versa has been confirmed for the artificial exercise tests only (tiptoe movements, walking in place etc.).
- The main calf muscle pump function was considered as its ability to eject blood centripetally by posterior calf muscle group contraction.

**What New Information Does This Article Contribute?:**

- An absence of pressure gradient directed from the calf to the thigh (centripetal) in the great the saphenous vein during locomotion.
- During locomotion, at the lower leg level, the primary route of blood outflow from the superficial veinous network toward intramuscular veins is a horizontal route (through perforating veins).
- New parameters are introduced to assess calf muscle pump effectiveness as its ability to maintain accordance between the muscle pump output and arterial blood supply during locomotion.

The calf muscle pump (CMP) is known as a significant contributor to venous blood outflow from the lower extremity due to its ability to effectively eject blood in a centripetal direction. Therefore, CMP failure refers to an impaired ejecting ability associated with chronic venous insufficiency (CVI) occurrence and progression. It is expressed in the increase in ambulatory venous pressure referred to as ambulatory venous hypertension (AVH). However, multiple studies demonstrated a lack of agreement between defined CMP failure and AVH, the severity of CVI, and quality of life.

Here we show that during locomotion, the CMP acts as a stream diversion pump redirecting blood flow from superficial veins (SVs) to intramuscular veins (IVs) through perforating veins. This is because the observed pressure gradient prevents centripetal blood flow from the calf to thigh level in the SVs and favors blood flow from SVs to IVs during the swing phase of the stride cycle. This function is provided by the synergetic work of antagonist calf muscles (anterior tibial muscle and gastrocnemius). Thereby, the CMP prevents pressure growth in the superficial veins of the lower leg (AVH) during exercise when the arterial blood supply increases according to exercise intensity.

## Introduction

Chronic venous insufficiency (CVI) is a term reserved for advanced chronic venous disease (CVD) including edema, skin changes, and venous ulcers, and it affects 1 – 11% of the general population, with a significant socio-economic impact.^1–5^ There is a consensus that CVI is associated with ambulatory venous hypertension (AVH) caused by either venous flow abnormalities (reflux and/or obstruction) and/or extra-venous pathology, such as orthopedic disorders, obesity, etc. In a standing position, venous pressure is determined mainly by hydrostatic pressure, and at the lower leg and foot levels it ranges on average from 60 to 90 mmHg depending on the distance from the measurement point to the right atrium.^6–11^ In turn, during an exercise such as walking, tiptoe movements, knee bending, etc., venous pressure drops significantly.^7,10,12^ The drop in venous pressure during exercise is termed ambulatory venous pressure (AVP). AVP is usually measured in the dorsal foot (superficial) vein. Its maximum cut-off value is 30 mmHg. Exceeding values are defined as AVH. ^4,5,7,13–20^ AVP is the best discriminative parameter for estimating both hemodynamic abnormalities and CVD severity.^21^

Because the drop in venous pressure in the superficial venous network during exercise is attributed to CMP function, several studies have been conducted to establish the relationships between the functional tests assessing CMP efficiency (ejection fraction [EF] and residual volume fraction [RVF] of plethysmography), AVP, disease severity, and quality of life. The early studies reported a wide range of relationships between functional tests and AVP and/or disease severity from negligible ^22,23^ to moderate.^24–28^ However, recent studies have shown a weak relation of EF and RVF to either AVP or disease severity.^16,29^ Moreover, attempts to ameliorate CMP function using resistance exercise programs have shown a lack of agreement between clinical outcomes and the results of improved CMP functional parameters.^29–33^ The functional tests are based on the assumption that CMP effectiveness is the ability to eject blood from the largest intramuscular veins (venous sinuses [VS]) of the gastrocnemius (GCM) and soleus muscles by their contraction. In turn, to provide blood ejection, VS should be adequately filled with blood between muscle contractions (relaxation phase).^18,19^ However, a recent study assessing CMP work during actual ambulation has shown that the muscle pump function differs from artificial maneuvers in the functional tests.^34^ First, the VS acts as a conduit rather than a reservoir. It transfers blood from the network of intramuscular veins to the axial deep veins.

Second, the CMP is not a mechanism of the posterior calf muscle group only; instead, it includes the synergetic work of antagonist muscles and the ankle joint: GCM contraction with parallel ATM relaxation (generating high pressure in intramuscular veins [IV]) during the stance phase and, vice versa, ATM contraction with parallel GCM relaxation (generating very low or even negative IV pressure) during the swing phase of the gait cycle (**Figure 1 b, c**).

**Figure 1.**
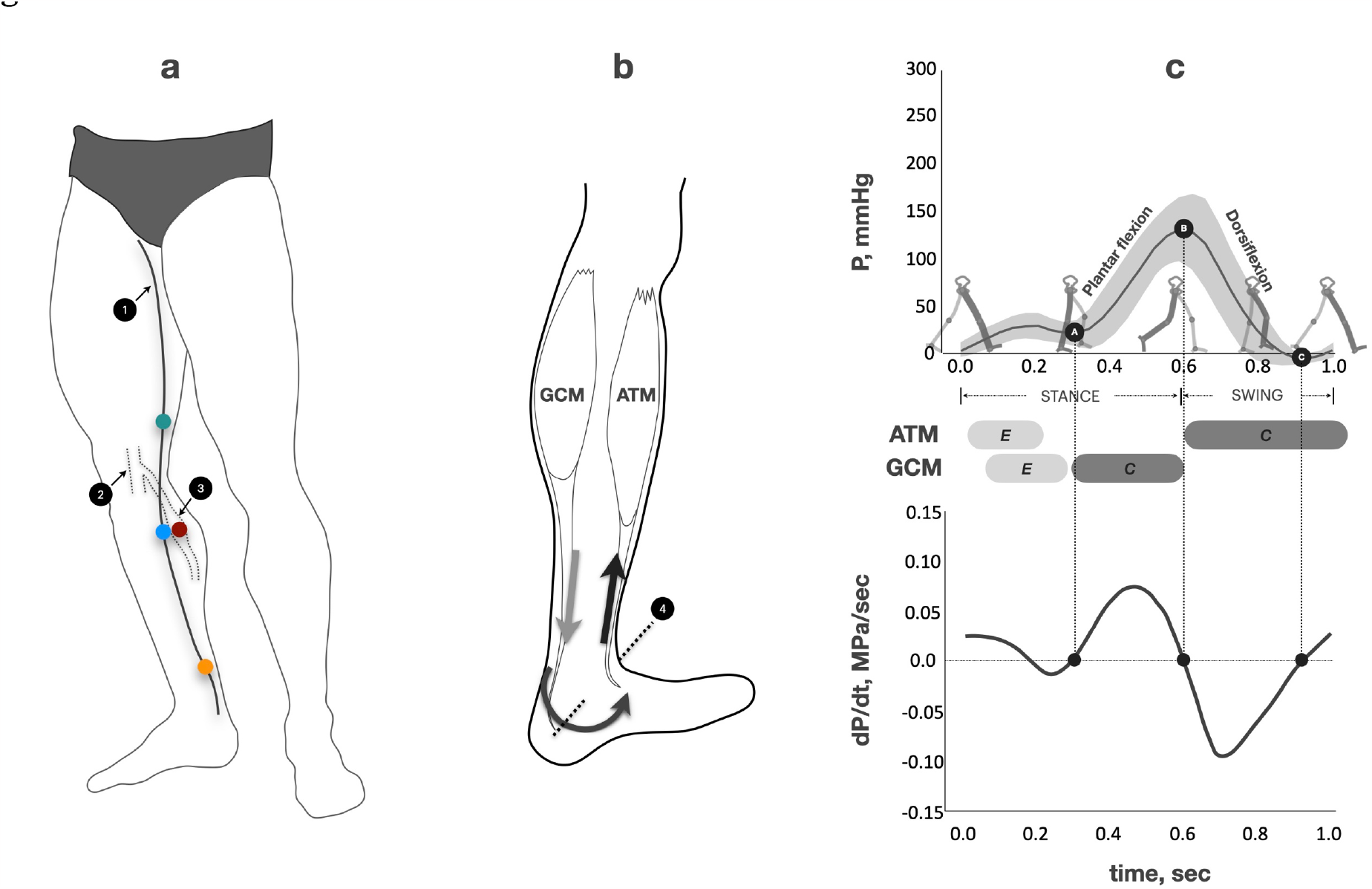
**Study methodology** **Figure 1a: *Pressure measurement*** ***1-*** great saphenous vein (GSV) ***2-*** popliteal vein ***3-*** intramuscular vein (IV) of gastrocnemius ***Orange point*** – GSV at ankle level ***Blue point*** – GSV at proximal calf level ***Green point*** – GSV at mid-distal thigh level ***Red point*** – IV at proximal calf level **Figure 1b: *Biomechanics of dorsiflexion*** ***4-*** axis of ankle joint rotation ***GCM*** – gastrocnemius muscle ***ATM*** – anterior tibial muscle ***Arrows*** – direction of movement **Figure 1c: *Methodology of calf muscle pump unit power calculation*** ***The top graph is the pressure changes in the intramuscular veins of the GCM across the gait cycle of ambulation***. ***The bottom graph is the pressure derivative (unit power of muscle pump***, 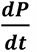***) changes across the gait cycle of ambulation***. ***A*** is the point when GCM begins concentric contraction (heel rise moment). ***B*** is the point of the highest pressure corresponding to the end of GCM contraction, and the following foot take-off when ATM starts its concentric contraction for dorsiflexion. ***C*** is the point of the lowest pressure approximately corresponding to the end of ATM concentric contraction (foot dorsiflexion). ***AB*** corresponds to the positive values of 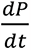 ***BC*** corresponds to the negative values of 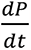 ***Dotted projection lines*** show that points **A, B**, and **C** correspond to zero value of the 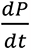 ***GCM*** – gastrocnemius muscle ***ATM*** – anterior tibial muscle ***Light grey boxes “E”*** – timeline of eccentric (increase in tension with muscle lengthening) muscle contraction ***Dark grey boxes “C”*** – timeline of concentric (increase in tension with muscle shortening) muscle contraction

We hypothesize that the change in venous pressure in superficial veins and its relatively low value (AVP ≤ 30 mmHg) during exercise is provided by the two consecutive phases of the CMP: a suction of blood from superficial veins to IVs, and centripetal blood ejection from IVs. Consequently, the CMP acts like a stream diversion pump redirecting blood flow from superficial to IVs during natural locomotion. CMP effectiveness during exercise is determined by the ability to maintain balance between the output of the muscle pump and arterial blood supply.

This study aimed to investigate the real-time relationship between GCM and ATM activity and pressure gradient (PG) changes at the different GSV segments and between the GSV and IV of the GCM during exercises with different types of biomechanics.

## Methods

The methods used were thoroughly described in the previous work.^34^ The study conformed to the Declaration of Helsinki. The local ethics committee of Surgut State University approved the study (protocol #22 of 02.17.2021). The same sample of nine healthy volunteers (seven men and two women) was enrolled in the study.^34^ None had clinical or ultrasound signs of CVD, impaired joint mobility, or artery lesions of the lower extremities. The participants were carefully informed of the study protocol and provided written consent to participate in the study. The subjects’ anthropometrics (mean ± SD) were age 32.8 ± 5 years, height 177 ± 5 cm, weight 79.7 ± 16 kg, body mass index (BMI) 25 ± 4 kg/m^2^, and calf circumference 38.2 ± 3.3 cm.

Subjects came to the laboratory at 10:00 AM and were instructed not to eat or drink coffee or tea and to avoid physical exercise for two hours before the experiment. Then, volunteers rested in a lying position for around 30 – 40 minutes while catheterization of the target veins was done under local anesthesia using ultrasound guidance. Three intravenous catheters 18 gauge filled with heparinized saline were placed in the GSV at the ankle [GSV_ANKLE_], upper third of the calf [GSV_CALF_], and mid-distal thigh [GSV_THIGH_] levels. A 20-cm-long catheter (*Balton Sp*.*z*.*o*.*o*., *Warszawa, Poland*) filled with heparinized saline was placed in the large intramuscular vein of the gastrocnemius medial aspect (IV_CALF_). The intramuscular vein approaching for catheterization was connected to the superficial venous system via a perforating vein (PV) with size and anatomy allowing for catheterization without muscle damage.^34^ The intramuscular catheter was adjusted to the level of the GSV_CALF_ catheter. All catheterization points are presented in **Figure 1a**. The catheters were secured by cannula retention dressing. Pressure transducers (*TruWave 3 cc/60 PX260, Edwards Lifescience Services GmbH, Germany*) connected to the catheters were secured on the lateral aspect of the calf and thigh at the same level as the catheters. A multi-channel blood pressure analysis system, “*Angioton*” (*BIOSOFT-M Ltd*., *Moscow, Russia*), was used for continuous pressure monitoring. Two electromyography (EMG) probes (*Callibri Group Ltd*., *Saint Petersburg, Russia*) were placed over the GCM medial aspect and ATM after the preparation for pressure measurement. The probes were connected to the system software installed on a personal computer by Bluetooth. The software displayed EMG data as real-time graphs.

Then, volunteers consistently completed the following exercise tests. First, treadmill (*Physioline TNX TOUCH, Svensson Body Labs, China*) walking (W) with a frequency of 30, 45, and 60 stride cycles min^-1^. The stride cycle is the time from the heel strike of one foot to the next heel strike by the same foot; during locomotion, the number of stride cycles is half of the gait cycles (steps).^35^ Second, treadmill running with 75 and 90 strides min^-1^. Third, plantar flexions in a standing position with 30, 45, and 60 flexions min^-1^. During the PF test, subjects stood on the treadmill while holding onto its frame. Every exercise bout lasted one minute and was separated by 10 minutes of rest. The laboratory room temperature was adjusted to the thermoneutral zone 21-23 °C.

The blood volume flow rate in the common femoral artery of the studied leg (LBF) was calculated as the product of the cross-sectional area (CSA) received from the ultrasound B-mode image, the time-average velocity (TAMEAN), and the heart rate (HR), both measured in a longitudinal plane and averaged over a 12-sec Doppler scan (LBF = CSA*TAMEAN*HR, liter/min). The ultrasound measurements were taken in a standing still position once before the exercise session and immediately after the end of each exercise bout. During exercise, the pressure and EMG data were recorded continuously. Each exercise set was recorded by a video camera embedded in a smartphone (*iPhone 11 Pro, Apple Inc*., *USA*) using 1080 pixels in a 60-frames-per-sec format for the ensuing evaluation of the stride (W and R) and PF cycles using frame-by-frame analysis of the videos.^36,37^

Because walking, running, and plantar flexion are cyclic processes, the time-pressure curves of 10 subsequent strides or flexions of each subject were reinterpolated to a uniform grid with constant time step and then averaged to either single stride or PF cycle for the ensuing data comparison. To avoid observer bias, the data analysis (the time of stride and PF cycle events, pressure, and EMG parameters) was performed using 10 consecutive strides (movements) in the middle of each exercise bout. The cycle time-averaged pressure (mean pressure 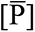, mmHg) was calculated for each GSV point for each exercise set. The pressure gradient (PG, mmHg) between adjacent points of measurement was calculated as *P*_*vein*1_ − *P*_*vein*2_ for each time moment of either the stride or PF cycle. Then, PG values were plotted on a graph for each type of exercise bout. In total, four PG curves were plotted: PG_RED_ (IV_CALF_ minus GSV_CALF_), PG_WHITE_ (IV_CALF_ minus GSV_ANKLE_), PG_ORANGE_ (GSV_CALF_ minus GSV_THIGH_), and PG_BLUE_ (GSV_ANKLE_ minus GSV_CALF_).

We analyze the rate of work per unit volume of blood which is associated with the change in kinetic energy of the blood flow and potential energy of the elastic stretching of veins per unit volume of blood and unit time. According to the basic principles of fluid dynamics ^38^, this value equals the pressure derivative by time

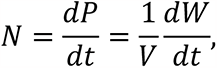

Where *N* is the unit power of the muscle pump, *P* is pressure, *V* is volume, and *W* is the work of the muscle pump, which is performed by the muscle pump to move a portion of the blood of volume V. For exercises with different frequencies, we compare the unit power of the muscle pump over the same time period *T*

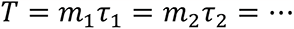

Where *m* is the number of strides or PF cycles and τ is the duration of a stride cycle in an exercise.

Thus, we compare the values 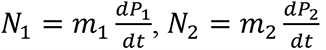, etc. The value of *N*_*j*_ equals the power of the muscle pump over period *T* in exercise *j*.

We compute the pressure derivate 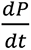 based on the measured pressure values *P* at time moments *t*_1_, *t*_2_, …,*t*_*N*_ (*N* is the total number of time moments) by the following finite difference methods

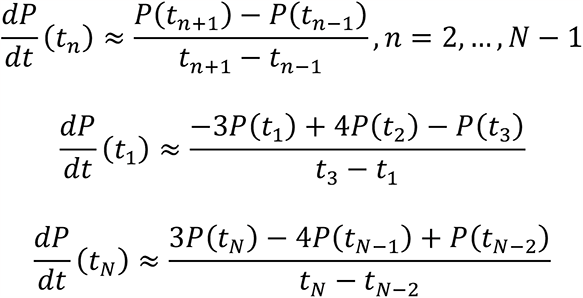

According to the theory of numerical methods, these formulas have the second order of accuracy.^39^

The calf muscle pump during locomotion is a cyclic process including two consecutive phases, which are ejection and suction with blood due to the work of antagonist calf muscles (GCM and ATM). ^34^ The force of GCM concentric contraction (an increase in tension with muscle shortening) generates blood ejection. In turn, the force of ATM concentric contraction with parallel GCM relaxation generates very low, even negative, pressure in the intramuscular venous network of the GCM (**Figure 1b, c**). The force of those muscles is reflected in the typical pressure curve of IV as segments AB and BC (**Figure 1c**), where “A” is the point at which the GCM begins concentric contraction (heel raise moment) and “B” is the point of the highest pressure corresponding to the end of GCM contraction and the following foot take-off when ATM starts its concentric contraction for dorsiflexion. “C” is the point of the lowest pressure approximately corresponding to the end of ATM concentric contraction (foot dorsiflexion). Points A and C correspond to the local minimum of *P*(*t*), while point B corresponds to the local maximum of *P*(*t*). In all measurements, we observe that AB is monotonically increasing part of *P*(*t*), while BC is monotonically decreasing part of *P*(*t*). According to the properties of smooth functions, AB corresponds to the positive values of 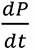, while BC corresponds to the negative values of 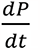. According to a necessary condition for an extremum of a smooth function, points A, B, and C correspond to zero value of 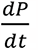. Thus, we identify *t*_*A*_, *t*_*B*_, *t*_*C*_ by setting the first derivative of *P*(*t*) to zero and finding corresponding values *t*_*A*_, *t*_*B*_, *t*_*C*_ with the finite difference methods shown above (see the lower row of graphs in **Figure 1c**). Thus, the work of GCM and ATM per cycle can be evaluated as time-averaged unit power for the periods AB (power of ejection, 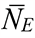) and BC (power of suction, 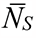), respectively. Finally, the average unit power per minute was calculated as 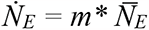 and 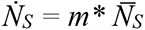 for each type of exercise, where 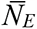 and 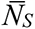 are cycle-averaged unit power of ejection and suction, respectively, and *m* is the number of either stride or flexion cycles per minute.

## Statistical analysis

Means and standard deviations were used to describe the continuous variables and their mathematical analysis. Nonparametric analyses were performed for statistical evaluation of the data (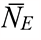, 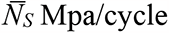, and 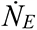 and 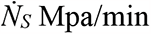, LBF L/min, and 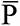 mmHg). The Friedman test and post hoc paired Mann-Whitney-Wilcoxon test with the Benjamini-Yekutieli correction were used to compare the parameters between different exercise frequencies. The Mann-Whitney-Wilcoxon test was used to compare the parameters between different types of exercise at the corresponding frequency. For multiple comparisons, the Benjamini-Yekutieli correction was used. The correlation analysis used the Spearman rank correlation coefficient. Statistical significance was defined as P < .05. The analysis was performed with Statistica 13 (*Statsoft Inc*., *Tulsa, Oklahoma, USA*), R Language 4.3 (*R Foundation, Vienna, Austria*), and Rstudio 2023.03 (*Posit Software, PBC, Boston, MA, USA*).

## Results

Venous pressure in all GSV points of measurement was different in the standing still position but it decreased to approximately the same value during exercise **(Figure 2)**. There was no significant difference in cycle time-averaged pressure (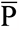, mmHg) between three GSV points at any frequency used during walking, running, or plantar flexion **(Figure 2b, c)**. In addition, there was no difference in 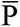 values at all GSV points between the used exercise frequencies. However, the 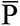 at each GSV point was statistically significantly higher during the PF exercise compared to the corresponding frequency of the walking exercise. **Figure 2d** shows that P averaged over three GSV points was 50% higher during plantar flexion than during walking regardless of exercise frequency.

**Figure 2.**
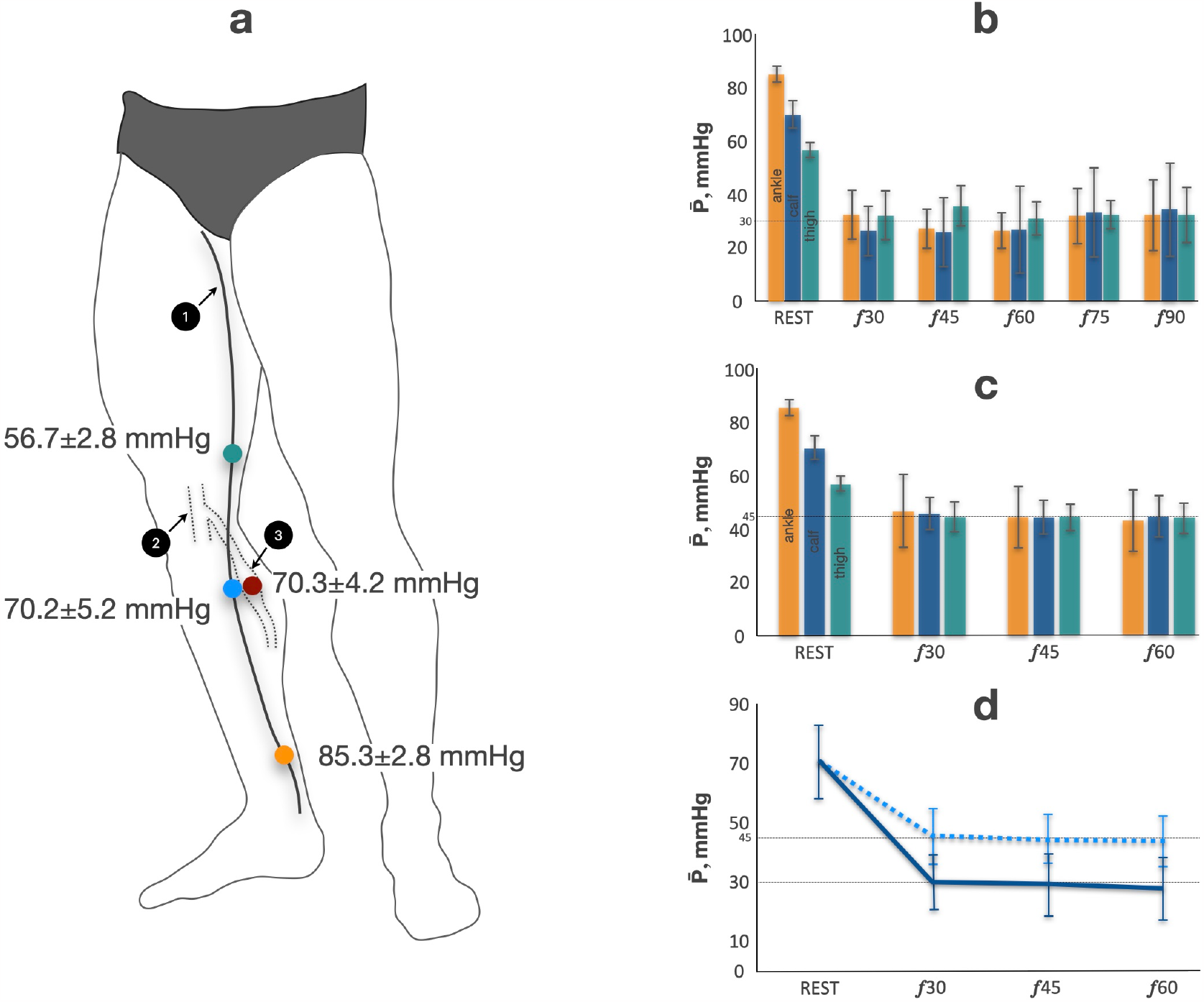
**Figure 2a: *Pressure data at standing still position*** ***1-*** great saphenous vein (GSV) ***2-*** Popliteal vein ***3-*** intramuscular vein (IV) of gastrocnemius (GCM) ***Orange point*** – GSV at ankle level ***Blue point*** – GSV at proximal calf level ***Green point*** – GSV at mid-distal thigh level ***Red point*** – IV at proximal calf level **Figure 2b: *Cycle time-average pressure during walking and running*** ***Orange column*** – cycle time-average pressure at GSV ankle level ***Blue column*** – cycle time-average pressure at GSV proximal calf level ***Green column*** – cycle time-average pressure at GSV mid-distal thigh level ***Rest*** – standing still position ***f*** – stride frequency per minute **Figure 2c: *Cycle time-average pressure during plantar flexion*** ***Orange column*** – cycle time-average pressure at GSV ankle level ***Blue column*** – cycle time-average pressure at GSV proximal calf level ***Green column*** – cycle time-average pressure at GSV mid-distal thigh level ***Rest*** – standing still position ***f*** – flexions frequency per minute **Figure 2d: *Cycle time-average pressure averaged by three GSV points*** ***Blue dotted line*** – cycle time-average pressure in GSV during plantar flexion ***Blue solid line*** – cycle time-average pressure in GSV during walking ***Rest*** – standing still position ***f*** – movement frequency per minute

The time-dependent pressure curves of stride (walking and running) and PF cycles are presented in **Figures 3, 4**, and **5** (upper graphs), respectively. The shape of the pressure curve was similar at all points IV_CALF_, GSV_CALF_, GSV_ANKLE_, and GSV_THIGH_ for particular types of exercise independent from the exercise frequency or studied subject. The amplitude of pressure in the intramuscular vein was substantially greater than the pressure amplitude in all GSV points for all exercises. The pressure amplitude in the GSV gradually decreases with the increase in stride or plantar flexion frequency. Thus, the curves of mean values are almost converged at 60, 75, and 90 strides or 45 and 60 flexions min^-1^. The observed shape of IV_CALF_ pressure was significantly different between the three types of exercise. There was no pressure “plateau” at the beginning of the stance phase during running, unlike with walking (**Figure 3**,**4**). The pressure of the PF cycle had two substantial consecutive peaks. The first peak corresponded to GCM concentric contraction (an increase in tension with muscle shortening) when the heel was raised (**Figure 5**). A second, even higher peak was observed when GCM eccentrically contracted (an increase in tension with muscle lengthening) while the heel was going down. When the heel had already been put down, the GCM continued contracting until the body returned to a posture perpendicular to the ground. The most pronounced change in pressure took place after a heel had already been grounded. The minimum IV_CALF_ pressure (P_min_) during PF cycles was significantly higher than the P_min_ of the corresponding stride (walking) cycles (PF_30_ 42.1±11.6 vs. W_30_ 1.9±3.8 mmHg, P < .0001; PF_45_ 46.1±16.5 vs. W_45_ 0.3±4.1 mmHg, P < .0001; PF_60_ 57.2±24.1 vs. W_60_ - 4.6±3.5 mmHg, P < .0001).

**Figure 3.**
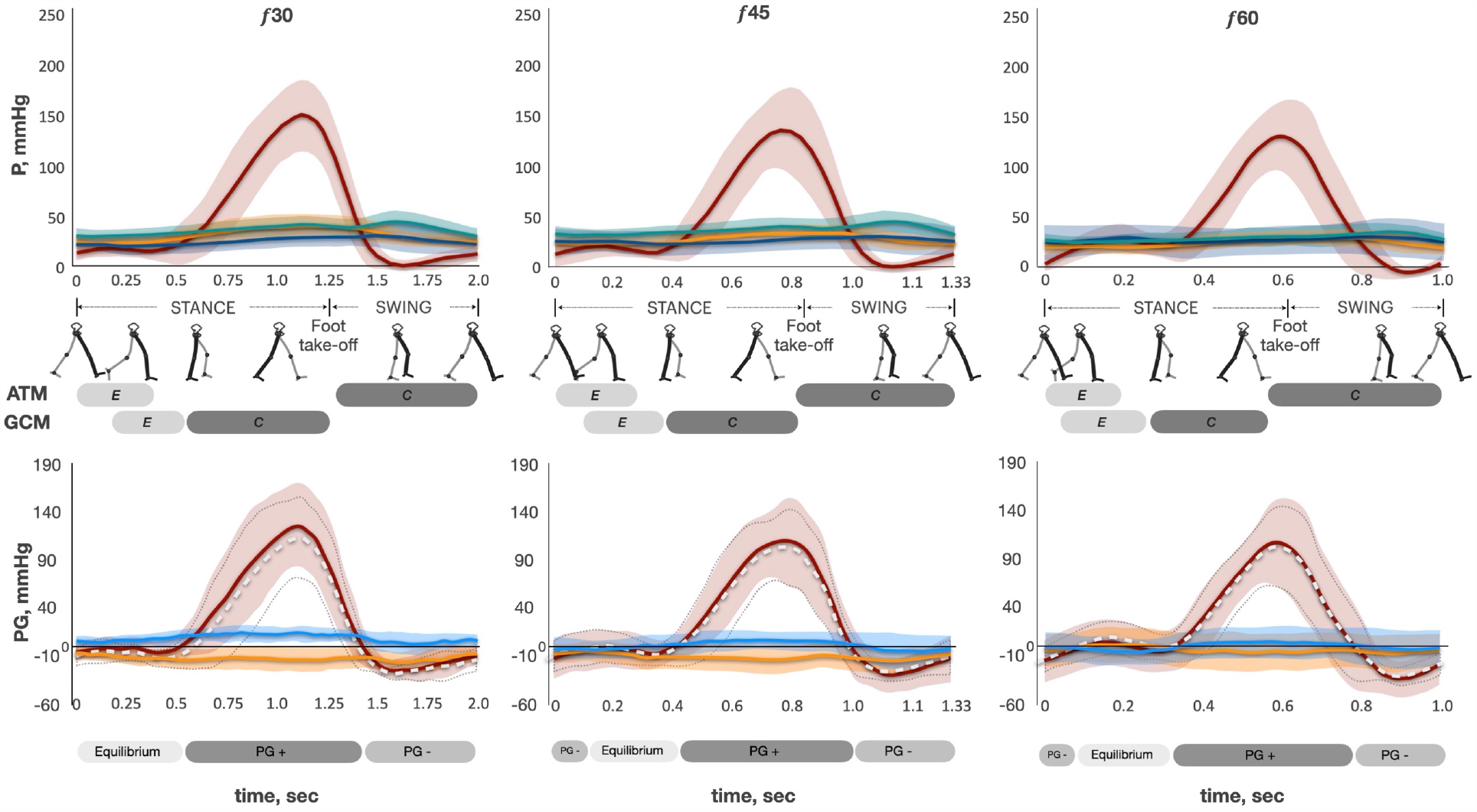
Legend: ***Pressure and pressure gradient changes across the stride cycle of ambulation. The top row is pressure curves at different stride cycle frequencies***. ***P*** – pressure ***Solid red curve*** – mean pressure of intramuscular vein of GCM (IV_CALF_) ***Light red area*** – standard deviation of mean IV_CALF_ pressure ***Solid green curve*** – mean pressure of GSV at mid-distal thigh level (GSV_THIGH_) ***Light green area*** – standard deviation of mean GSV_THIGH_ pressure ***Solid blue curve*** – mean pressure of GSV at proximal calf level (GSV_CALF_) ***Light blue area*** – standard deviation of mean GSV_CALF_ pressure ***Solid orange curve*** – mean pressure of GSV at ankle level (GSV_ANKLE_) ***Light orange area*** – standard deviation of mean GSV_ANKLE_ pressure ***f*** – stride frequency per minute ***ATM*** – anterior tibial muscle ***GCM*** – gastrocnemius muscle ***Light grey boxes “E”*** – timeline of eccentric (increase in tension with muscle lengthening) muscle contraction ***Dark grey boxes “C”*** – timeline of concentric (increase in tension with muscle shortening) muscle contraction ***The bottom row is pressure gradient curves at different stride cycle frequencies***. ***PG*** – pressure gradient ***Solid red curve*** – mean of PG between IV_CALF_ and GSV_CALF_ (PG_RED_ = P IV_CALF_ – P GSV_CALF_) ***Light red area*** – standard deviation of PG_RED_ ***Dotted white curve*** – mean of PG between IV_CALF_ and GSV_ANKLE_ (PG_WHITE_ = P IV_CALF_ – P GSV_ANKLE_) ***Area bounded by dotted grey line*** – standard deviation of PG_WHITE_ ***Solid blue curve*** – mean pressure of PG between GSV_ANKLE_ and GSV_CALF_ (PG_BLUE_ = P GSV_ANKLE_ – P GSV_CALF_) ***Light blue area*** – standard deviation of PG_BLUE_ ***Solid orange curve*** – mean pressure of PG between GSV_CALF_ and GSV_THIGH_ (PG_ORANGE_ = P GSV_CALF_ – P GSV_THIGH_) ***Light orange area*** – standard deviation of PG_ORANGE_ ***Light boxes “Equilibrium”*** – timeline of negligible PG between IV and GSV ***Light grey boxes “PG -”*** – timeline of negative PG directed from GSV to IV ***Dark grey boxes “PG +”*** – timeline of positive PG directed from IV to GSV

**Figure 4.**
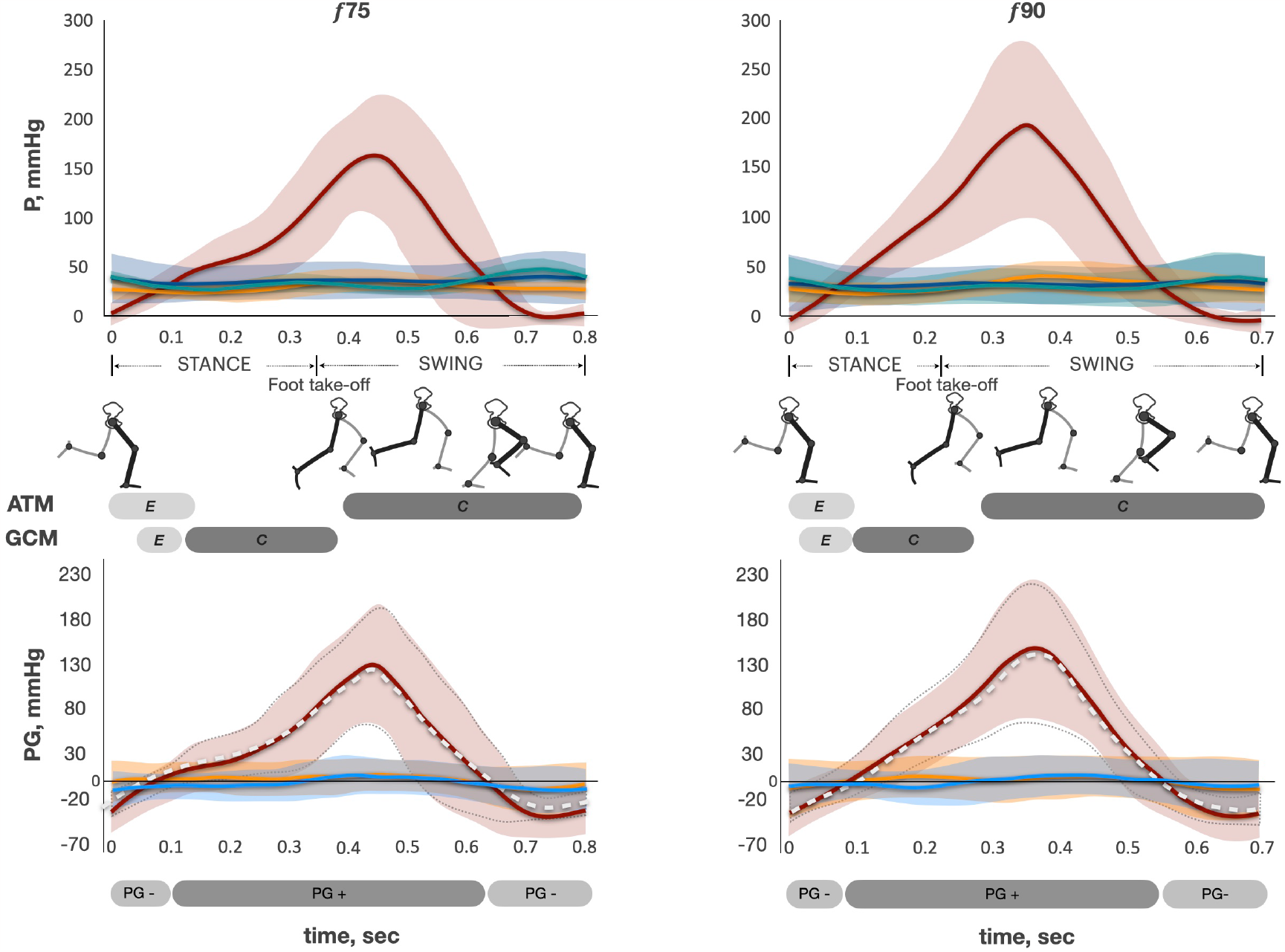
Legend: ***Pressure and pressure gradient changes across the stride cycle of running. The top row is pressure curves at different stride cycle frequencies***. ***P*** – pressure ***Solid red curve*** – mean pressure of intramuscular vein of GCM (IV_CALF_) ***Light red area*** – standard deviation of mean IV_CALF_ pressure ***Solid green curve*** – mean pressure of GSV at mid-distal thigh level (GSV_THIGH_) ***Light green area*** – standard deviation of mean GSV_THIGH_ pressure ***Solid blue curve*** – mean pressure of GSV at proximal calf level (GSV_CALF_) ***Light blue area*** – standard deviation of mean GSV_CALF_ pressure ***Solid orange curve*** – mean pressure of GSV at ankle level (GSV_ANKLE_) ***Light orange area*** – standard deviation of mean GSV_ANKLE_ pressure ***f*** – stride frequency per minute ***ATM*** – anterior tibial muscle ***GCM*** – gastrocnemius muscle ***Light grey boxes “E”*** – timeline of eccentric (increase in tension with muscle lengthening) muscle contraction ***Dark grey boxes “C”*** – timeline of concentric (increase in tension with muscle shortening) muscle contraction ***The bottom row is pressure gradient curves at different stride cycle frequencies. PG*** – pressure gradient ***Solid red curve*** – mean of PG between IV_CALF_ and GSV_CALF_ (PG_RED_ = P IV_CALF_ – P GSV_CALF_) ***Light red area*** – standard deviation of PG_RED_ ***Dotted white curve*** – mean of PG between IV_CALF_ and GSV_ANKLE_ (PG_WHITE_ = P IV_CALF_ – P GSV_ANKLE_) ***Area bounded by dotted grey line*** – standard deviation of PG_WHITE_ ***Solid blue curve*** – mean pressure of PG between GSV_ANKLE_ and GSV_CALF_ (PG_BLUE_ = P GSV_ANKLE_ – P GSV_CALF_) ***Light blue area*** – standard deviation of PG_BLUE_ ***Solid orange curve*** – mean pressure of PG between GSV_CALF_ and GSV_THIGH_ (PG_ORANGE_ = P GSV_CALF_ – P GSV_THIGH_) ***Light orange area*** – standard deviation of PG_ORANGE_ ***Light grey boxes “PG -”*** – timeline of negative PG directed from GSV to IV ***Dark grey boxes “PG +”*** – timeline of positive PG directed from IV to GSV

**Figure 5.**
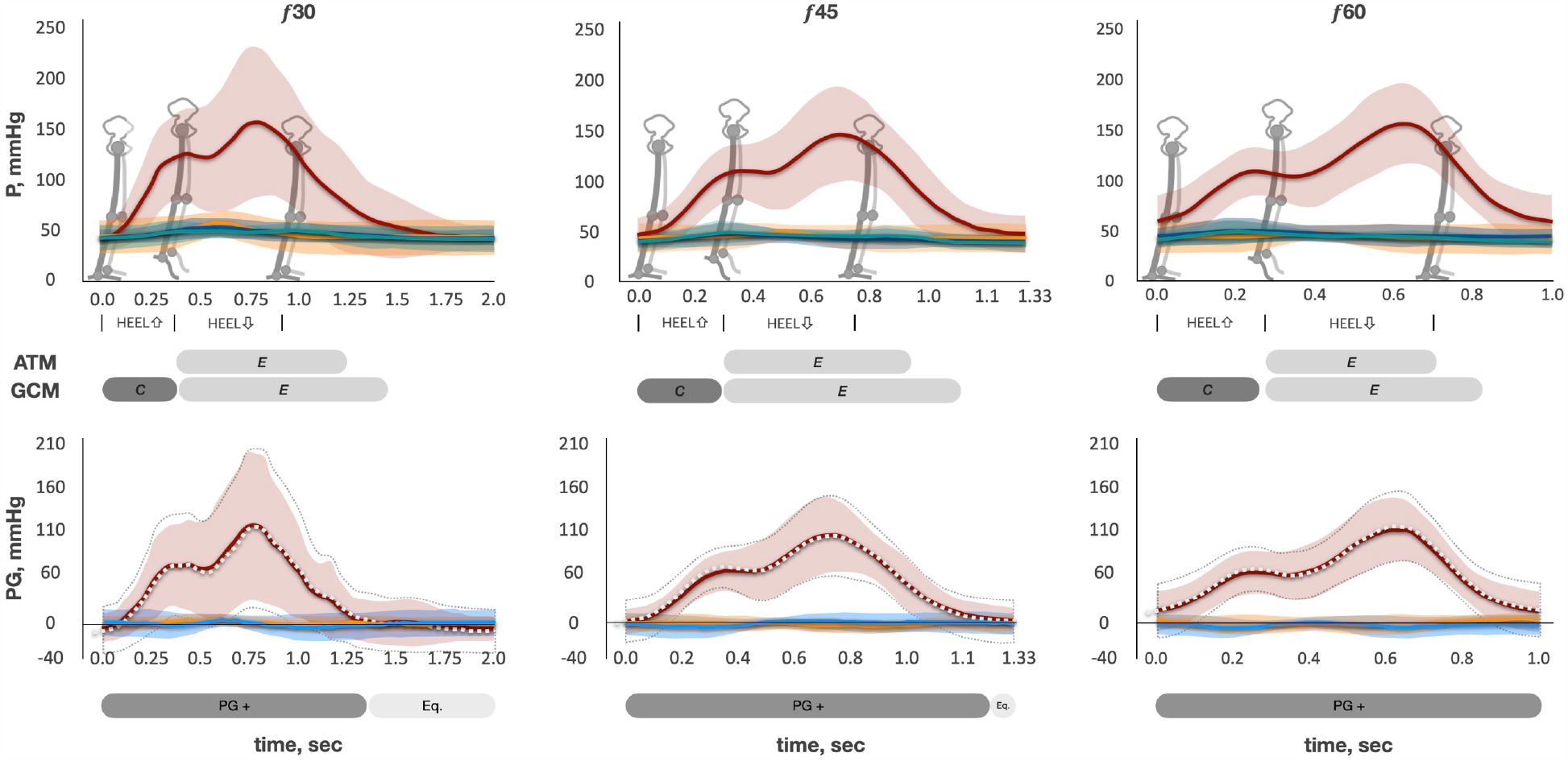
Legend: ***The pressure and pressure gradient changes across the plantar flexion cycle. The top row is pressure curves at different stride cycle frequencies***. ***P*** – pressure ***Solid red curve*** – mean pressure of intramuscular vein of GCM (IV_CALF_) ***Light red area*** – standard deviation of mean IV_CALF_ pressure ***Solid green curve*** – mean pressure of GSV at mid-distal thigh level (GSV_THIGH_) ***Light green area*** – standard deviation of mean GSV_THIGH_ pressure ***Solid blue curve*** – mean pressure of GSV at proximal calf level (GSV_CALF_) ***Light blue area*** – standard deviation of mean GSV_CALF_ pressure ***Solid orange curve*** – mean pressure of GSV at ankle level (GSV_ANKLE_) ***Light orange area*** – standard deviation of mean GSV_ANKLE_ pressure ***f*** – stride frequency per minute ***ATM*** – anterior tibial muscle ***GCM*** – gastrocnemius muscle ***Light grey boxes “E”*** – timeline of eccentric (increase in tension with muscle lengthening) muscle contraction ***Dark grey boxes “C”*** – timeline of concentric (increase in tension with muscle shortening) muscle contraction ***The bottom row is pressure gradient curves at different stride cycle frequencies. PG*** – pressure gradient ***Solid red curve*** – mean of PG between IV_CALF_ and GSV_CALF_ (PG_RED_ = P IV_CALF_ – P GSV_CALF_) ***Light red area*** – standard deviation of PG_RED_ ***Dotted white curve*** – mean of PG between IV_CALF_ and GSV_ANKLE_ (PG_WHITE_ = P IV_CALF_ – P GSV_ANKLE_) ***Area bounded by dotted grey line*** – standard deviation of PG_WHITE_ ***Solid blue curve*** – mean pressure of PG between GSV_ANKLE_ and GSV_CALF_ (PG_BLUE_ = P GSV_ANKLE_ – P GSV_CALF_) ***Light blue area*** – standard deviation of PG_BLUE_ ***Solid orange curve*** – mean pressure of PG between GSV_CALF_ and GSV_THIGH_ (PG_ORANGE_ = P GSV_CALF_ – P GSV_THIGH_) ***Light orange area*** – standard deviation of PG_ORANGE_ ***Light boxes “Equilibrium”*** – timeline of negligible PG between IV and GSV ***Dark grey boxes “PG +”*** – timeline of positive PG directed from IV to GSV

We analyzed the pressure difference (PG) between the pairs of adjacent points of measurement throughout the stride and PF cycles (see lower graphs in **Figures 3, 4**, and **5**). The PG_RED_ curve represents the changes in the pressure difference between IV_CALF_ and GSV_CALF_, mmHg. PG_WHITE_ represents the changes in the pressure difference between IV_CALF_ and GSV_ANKLE_, mmHg. PG_RED_ and PG_WHITE_ have similar shapes during walking, running, and PF exercise. The difference between PG_RED_ and PG_WHITE_ decreases with an increase in exercise frequency. Walking at 30 strides min^-1^ (**Figures 3**): During the first half of the stance phase (before heel rise), there was no significant PG between IV and GSV (curves are close to zero-line, “equilibrium phase”). Then, when the heel starts rising and GCM concentric contraction begins, PG rapidly grows (PG is directed from IV to GSV, “PG-positive phase”). The highest PG was observed by the end of GCM concentric contraction (end of stance phase). PG starts sharply decreasing at the beginning of the swing phase (the moment of foot take-off) when ATM initiates its concentric contraction and the GCM is already relaxed. It rapidly becomes negative (PG is directed from GSV to IV, “PG-negative phase”) and reaches its minimum value around the middle of the swing phase. Then, in the remaining swing phase, PG slowly comes to the zero-line. Further, an increase in the walking frequency (45 and 60 strides min^-1^) was characterized by a slightly higher PG change around the zero-line during the equilibrium phase. Because stride cycle duration decreases (*f* 30 2.0±0.04 sec, *f* 45 1.33±0.02 sec, *f* 60 1.0±0.01 sec, *f* 75 0.8±0.01 sec, and *f* 90 0.66±0.01 sec) with an increase in its frequency, the absolute time of all three phases decreases as well (**Figure 3**,**4** and **5**). A feature of running is the absence of the equilibrium phase (**Figure 4**). During the PF test, there was no PG-negative phase; the equilibrium phase was short at 45 flexions min^-1^ and absent at 60 flexions min^-1^ (**Figure 5**).

The PG_BLUE_ curve (pressure difference between GSV_ANKLE_ and GSV_CALF_, mmHg) always has positive values during the stride cycle at 30 strides min^-1^. This means that PG is directed centripetally from the ankle to the calf. The curve approaches the zero-line with the increase in frequency (45, 60, 75, and 90 strides min^-1^). The PG_ORANGE_ curve (P GSV_CALF_ minus P GSV_THIGH_, mmHg) always has negative values in the stride cycle at 30 strides min^-1^. This means that PG is directed retrogradely from the thigh to the calf. The curve gradually gets closer to zero with the increase in stride frequency (45 and 60 strides min^-1^) and becomes almost zero at 75 and 90 strides min^-1^.

Figure 6. demonstrates the changes in the average unit power of muscle pump suction 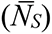 and ejection 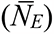 generated by a single stride or PF cycle. 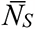 and 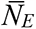 statistically significantly linearly increased with the increase in stride frequency (**Figure 6a)**. Both parameters remained unchanged with the increase in PF frequency (**Figure 6b**). The parameters were statistically significantly higher during walking compared to plantar flexion for the corresponding frequencies (**Figure 6 a, b**).

**Figure 6.**
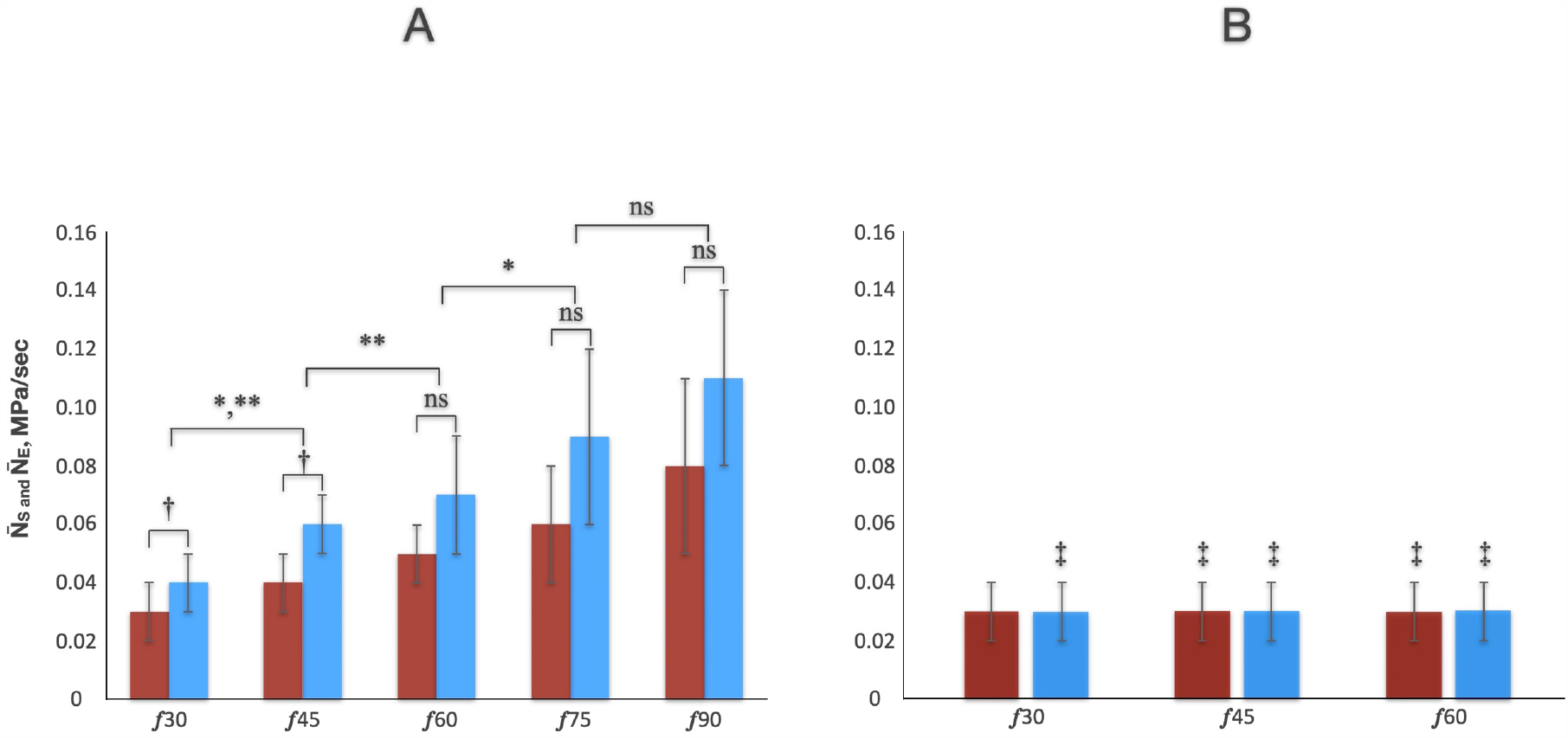
***A: Average unit power of calf muscle pump generated by single stride cycle*** ***B: Average unit power of calf muscle pump generated by single plantar flexion cycle Red column*** – cycle-averaged unit power of muscle pump ejection 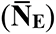 ***Blue column*** – cycle-averaged unit power of muscle pump suction 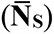 ***f*** – frequency per minute * – statistically significant difference between 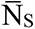 at different stride frequencies (P < 0.05, Friedman test and post hoc paired Mann-Whitney-Wilcoxon with Benjamini-Yekutieli correction for the multiple comparisons, f45 vs. f30, f60 vs. f45, f75 vs. f60, f90 vs. f 75) ** – statistically significant difference between 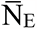 at different stride frequencies (P < 0.05, Friedman test and post hoc paired Mann-Whitney-Wilcoxon with Benjamini-Yekutieli correction for the multiple comparisons, f45 vs. f30, f60 vs. f45, f75 vs. f60, f90 vs. f 75) **†** – statistically significant difference between 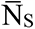 and 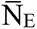 at corresponding frequencies (P < 0.05, Mann-Whitney-Wilcoxon test with Benjamini-Yekutieli correction for the multiple comparisons) ‡ – statistically significant difference between the parameters obtained during walking and plantar flexion exercises at corresponding frequencies (P < 0.05, Mann-Whitney-Wilcoxon test with Benjamini-Yekutieli correction for the multiple comparisons, f30 vs. f30, f45 vs. f45, and f60 vs. f60) ***ns*** – not significant (P > 0.05)

Arterial inflow (LBF, liter/min) and average unit power per minute (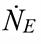 and 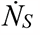, Mpa/min) are presented in **Figure 7**. LBF, 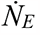, and 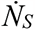 demonstrated similar exponential growth with the increase in stride frequency during walking and running. Exponential fitting of the functions *LBF*(*f*), 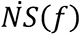, and 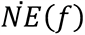 using the least squares method (LSM) produced the following results

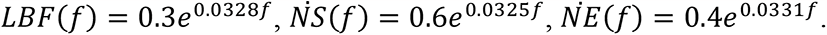

**Figure 7.**
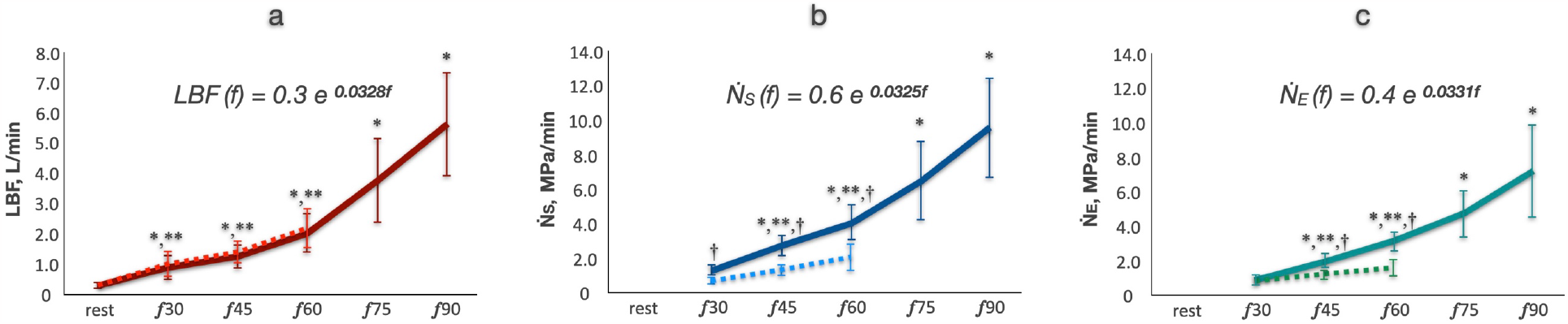
***A: Changes in volume flow rate in the common femoral artery (LBF) Solid red line*** – LBF changes during walking and running tests ***Dotted red line*** – LBF changes during plantar flexion tests ***B: Changes in minute unit power of calf muscle pump suction*** 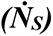 ***Solid blue line*** – 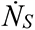 changes during walking and running tests ***Dotted blue line*** – 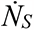 changes during plantar flexion tests ***C: Changes in minute unit power of calf muscle pump ejection*** 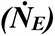 ***Solid green line*** – 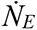 changes during walking and running tests ***Dotted green line*** – 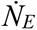 changes during plantar flexion tests ***f*** – frequency per minute * – statistically significant difference between the parameters (LBF, 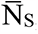, and 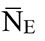) at different stride frequencies (P < 0.05, Friedman test and post hoc paired Mann-Whitney-Wilcoxon with Benjamini-Yekutieli correction for multiple comparison, f45 vs. f30, f60 vs. f45, f75 vs. f60, f90 vs. f 75) ** – statistically significant difference between the parameters (LBF, 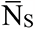 and 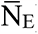) at different plantar flexion frequencies (P < 0.05, Friedman test and post hoc paired Mann-Whitney-Wilcoxon with Benjamini-Yekutieli correction for multiple comparison, f45 vs. f30, f60 vs. f45, f75 vs. f60, f90 vs. f 75) **†** – statistically significant difference between the parameters (LBF, 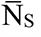, and 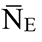) obtained during walking and plantar flexion exercises at corresponding frequencies (P < 0.05, Mann-Whitney-Wilcoxon test with Benjamini-Yekutieli correction for the multiple comparisons, f30 vs. f30, f45 vs. f45, and f60 vs. f60)

We observe that the power coefficients are very close in all cases. Thus, we conclude that 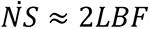 and 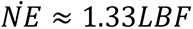 for all values of frequency *f*. The LSM and input errors explain the difference between the power coefficients. The strong connection between the parameters was confirmed by correlation analysis (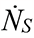 vs. LBF r = 0.91, 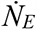 vs. LBF r = 0.9, P < .0001). The increase in LBF was comparable during PF and walking exercises. The increase in 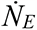 and 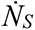 was significantly lower.

## Discussion

The main finding of the present study is the establishment of a pattern of pressure correlation between the intramuscular vein of the GCM and GSV, as well as between different levels of the GSV during locomotion and its changes in relation to the biomechanics of movement. We describe this pattern by the pressure gradient (PG, mmHg), which changes throughout the stride and plantar flexion cycles. The pattern has the following features. First is an absence of PG directed from the calf to the thigh (centripetal) in the GSV. Instead, a retrograde PG in the GSV was observed during all times of the stride cycle, but its magnitude decreased with the increase in the frequency of stride cycles (**Figure 3**,**4**). Second, a horizontal PG was observed between intramuscular and superficial veins. Its direction was determined by the joint work of antagonist calf muscles. Its value was considerably higher than the value of PG between any of the GSV levels (**Figure 3**,**4**). We introduce new parameters assessing CMP effectiveness, which are unit power of muscle pump ejection and suction. These parameters for the single stride cycle (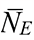 and 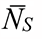) increased linearly with the increase in stride frequency (**Figure 6**). The average minute unit power of the muscle pump (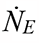 and 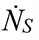) increased exponentially with the increase in stride frequency and was tightly coupled with the increased arterial blood supply of the lower extremity (**Figure 7**).

It was shown that blood ejection from the GCM during the stride cycle of ambulation takes place in a short period corresponding to GCM concentric contraction (from heel rise to foot take-off moment, that is, around 30% of stride cycle duration regardless of its frequency). The source reservoir is the intramuscular venous network except for the largest intramuscular veins (venous sinuses) that serve as a conduit transferring blood to the axial deep veins during ejection. At the beginning of the swing phase (foot take-off), ATM concentric contraction initiates active elongation of relaxed GCM (dorsiflexion, **Figure 1b, c**). It produces low or even negative pressure in the intramuscular veins of the GCM.^34^ Arterial flow to the working muscles occurs during their relaxation. ^40,41^ However, hypothetically, during muscle relaxation corresponding to the swing phase of the stride cycle, the intramuscular venous network should be filled with blood from the venous network located outside the muscle fascial sheath as well. This was shown by Almen and Nylander^18^ and Ludbrook^19^ in the case of plantar flexion with a 2-6 sec pause between contractions. We observed that PG is directed from either of the GSV levels to IV (PG-negative) from the middle swing phase until the loading response phase of the next stance. Thus, we confirm the same hypothesis for ambulation as well. We also note that PG in the GSV was directed from the thigh to calf level throughout the entire stride cycle for walking and running.

Thus, the probability of the occurrence of centripetal flow in the GSV is negligible at least from the calf to thigh level. We suppose that during locomotion the major route of blood flow from the superficial venous network toward the intramuscular veins runs through perforating veins (PVs). The absolute time of the PG-negative phase progressively decreases with the increase in stride frequency, while arterial blood flow increases exponentially (**Figure 7**). Most of the blood in the CFA is directed to working muscles. ^42,43^ The skin and subcutaneous blood flow of the exercising leg progressively increase as well.^44^ In this case, the pressure in the superficial veins should have risen along with the increase in the intensity of the exercise. On the contrary, we observe that the cycle time-averaged pressure at all levels of the GSV was stable with the increase in stride frequency (**Figure 2**). A similar pattern of the time-averaged pressure changes in the GSV at the ankle level was previously shown by Stick et al.^45^ and then Eifell et al.^46^ during treadmill walking and running tests. The possible explanation for the observed phenomenon is the increase in CMP productivity, which compensates for both the decreased blood flow time associated with the decreased PG-negative time and the increased arterial inflow. The increased CMP effectiveness came at the expense of both the increased cycle-averaged unit power of the muscle pump (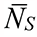 and 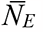, **Figure 6**) and the increase in stride frequency (average minute unit power of muscle pump, 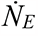, and 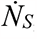, **Figure 7**). The latter parameters were tightly coupled with arterial blood inflow (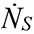 vs. LBF r = 0.91, 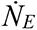 vs. LBF r = 0.9, P < .0001).

It can be speculated that hyperemic centripetal outflow from the superficial venous network through two main axial trunks (GSV and small saphenous vein) requires excessive energy expenses due to high hydraulic resistance. The hydraulic resistance of the vessel is inversely proportional to the vessel’s diameter in the fourth power and proportional to its lengths. Anatomical studies have shown that the skin and subcutaneous adipose tissue of almost all regions of the human lower extremity receive blood by the vessels going through or between muscle bellies (musculacutaneus and septocutaneous vessels, respectively). ^47–49^ The number of these perforating vessels is huge. There are more than 300 perforating arteries with a diameter greater than 0.5 mm on the lateral aspect of the lower leg.^49^ Every artery is accompanied by a couple of veins, so there is a huge number of perforating veins (PVs) linking the superficial venous network and the deep (intramuscular and intermuscular) veins at the lower leg level. While the diameter of a single PV varies from 0.5 mm to 2-3 mm, which is considerably less than the diameters of the GSV and SSV, the total diameter of PVs is significantly higher.

According to Kirchhoff’s law, the total effective resistance of several identical parallel conduits equals the resistance of one conduit divided by the number of conduits. In addition, the length of the PV is much smaller than the lengths of the GSV or SSV. Thus, the total effective hydraulic resistance of the network of PVs is rather small.

We conclude that during natural locomotion (walking and running), the CMP acts like a stream diversion pump redirecting blood flow from the superficial to intramuscular venous network through PVs (horizontal route) during the swing phase (ATM concentric contraction and GCM relaxation) and then ejecting blood to the axial deep veins (centripetally) during the second half of the stance phase (GCM concentric contraction) (**Figure 8**). We suggest that during locomotion, the horizontal route is likely the primary route of blood outflow from the superficial veins, at least at the lower leg level. The effective (optimal) CMP function means that the average minute unit power of muscle pump suction and ejection is proportional to the increased arterial blood supply during locomotion, that prevents the rise of pressure in superficial veins (AVP).

**Figure 8.**
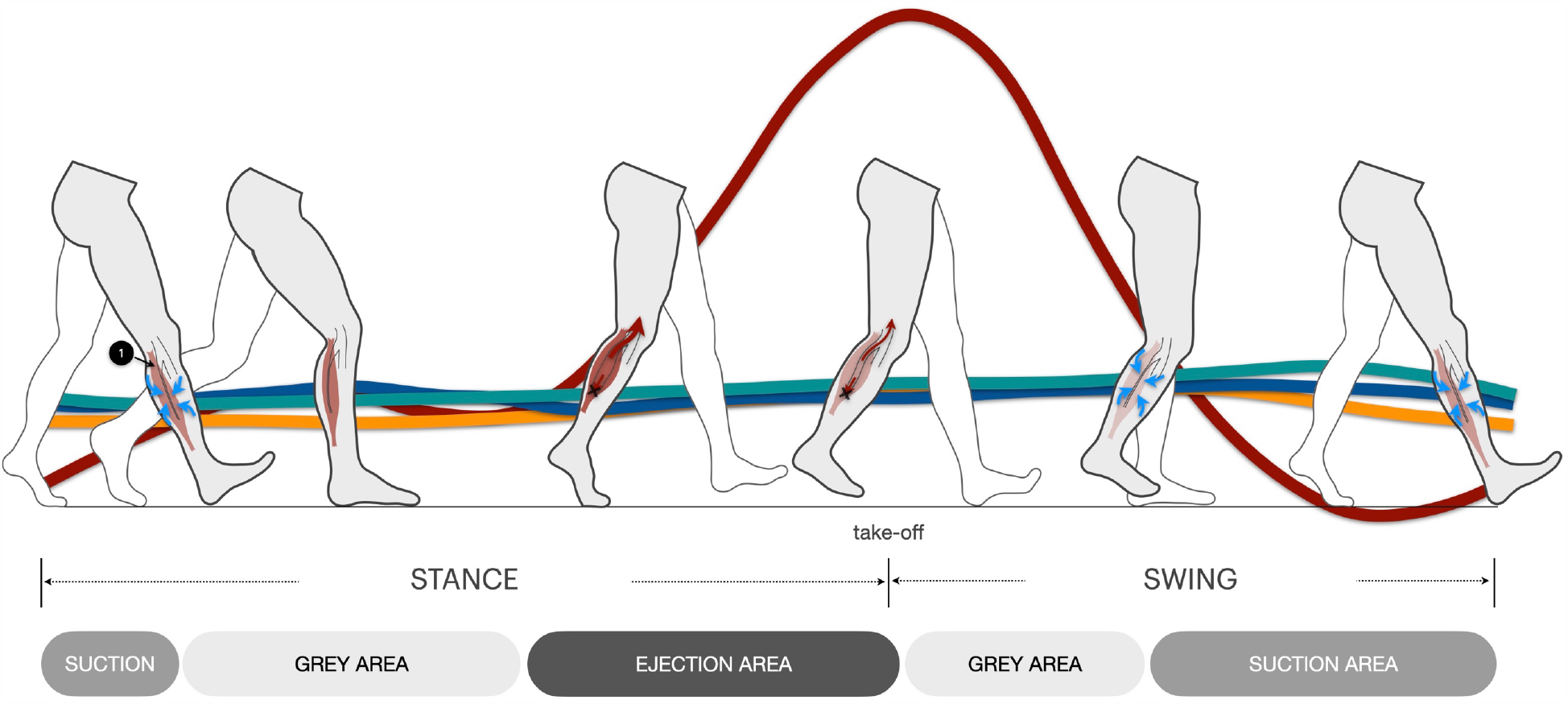
Legend: **1** – gastrocnemius muscle (GCM), the extent of red color saturation designates the extent of blood filling ***Solid red curve*** – pressure in intramuscular vein of GCM ***Solid green curve*** – pressure in great saphenous vein (GSV) at mid-distal thigh level ***Solid blue curve*** – pressure in GSV at proximal calf level ***Solid orange curve*** – pressure in GSV at ankle level ***Red arrows*** – direction of flow (centripetal) during GCM concentric contraction (second half of stance phase) ***Black cross*** – venous valves block blood flow ***Blue arrows*** – direction of flow toward intramuscular venous network of GCM ***Ejection area*** – timeline of blood ejection ***Suction area*** – timeline of blood flow from superficial to intramuscular veins ***Grey area*** – no significant blood exchange between intramuscular and superficial veins

It follows that impaired CMP productivity should lead to the deterioration of blood outflow resulting in increased AVP. While this suggestion has not been directly tested, the current study showed it by using another type of movement (plantar flexion) that does not have a phase with synergistic work of the antagonist calf muscles (ATM concentric contraction with parallel GCM relaxation). Instead, both muscles eccentrically contract to stabilize posture while the heel is going down. Therefore, the PF cycle-averaged unit power of the muscle pump (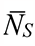 and 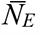, **Figure 6b**) was unchanged with the increase in flexion frequency. Meanwhile, the average minute unit power of the muscle pump demonstrated little increase that was uncoupled with LBF (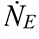, and 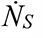, **Figure 7**). As a result, the cycle time-averaged pressure in the GSV was 50% higher during the PF exercises than during the corresponding walking frequencies. Such a difference is confirmed by the data from previous studies using similar exercise tests (treadmill walking and plantar flexion) for AVP measurement.^11,46,50^ Kügler et al.^50^ previously demonstrated that a lower degree of calf muscle activity by restricted ankle joint mobility increases AVP. In turn, it is well known that impaired ankle joint mobility correlates with advanced stages of chronic venous insufficiency. ^51–55^

### Study limitations

The main limitation of the study is the relatively small sample size. However, a strong consistency among the pressure data was observed in the studied limbs across different frequencies of used movement cycles for each exercise type. Technical challenges related to measuring blood flow volume in the CFA during exercise by ultrasound led the study team to use the LBF during postexercise hyperemia for correlation with the average minute unit power of the muscle pump. It has been shown that there is no difference between the LBF during exercise vs. an immediate postexercise LBF.^41^

## Conclusions

During the stride cycle of locomotion (walking and running), the CMP created two types of significant pressure gradients between intramuscular veins (IV) and superficial (GSV) veins (horizontal PG). First was PG-positive directed from the IV to the GSV owing to GCM concentric contraction (second half of the stance phase). Second was PG-negative directed from the GSV to the IV due to ATM concentric contraction with parallel GCM relaxation (swing phase). To evaluate the work performed by the GCM and ATM, we introduced the new parameters, which are the average minute unit power of suction 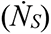 and ejection 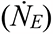. We found that these parameters are strongly correlated with changes in the arterial blood supply of the lower extremity during exercise tests.

## Nonstandard Abbreviations and Acronyms

ATM: anterior tibial muscle
AVP: ambulatory venous pressure
CFA: common femoral artery
CMP: calf muscle pump
CVD: chronic venous disease
CVI: chronic venous insufficiency
EF: ejection fraction
EMG: electromyography
GCM: gastrocnemius muscle
GSV: great saphenous vein
IV: intramuscular vein
LBF: blood volume flow rate in common femoral artery
PG: pressure gradient
RVF: residual volume fraction

## Acknowledgments

We are pleasure to appreciate Dr. Vladimir Denisov for his help in the study funding and organizational issues.

